# Calcineurin broadly regulates the initiation of skeletal muscle-specific gene expression by binding target promoters and facilitating the interaction of the SWI/SNF chromatin remodeling enzyme

**DOI:** 10.1101/628768

**Authors:** Hanna Witwicka, Jumpei Nogami, Sabriya A. Syed, Kazumitsu Maehara, Teresita Padilla-Benavides, Yasuyuki Ohkawa, Anthony N. Imbalzano

## Abstract

Calcineurin (Cn) is a calcium-activated serine/threonine protein phosphatase that is broadly implicated in diverse cellular processes, including the regulation of gene expression. During skeletal muscle differentiation, Cn activates the NFAT transcription factor but also promotes differentiation by counteracting the negative influences of protein kinase C beta (PKCβ) via dephosphorylation and activation of BRG1, an enzymatic subunit of the mammalian SWI/SNF ATP-dependent chromatin remodeling enzyme. Here we identified four major temporal patterns of Cn-dependent gene expression in differentiating myoblasts and determined that Cn is broadly required for the activation of the myogenic gene expression program. Mechanistically, Cn promotes gene expression through direct binding to myogenic promoter sequences and facilitating the binding of BRG1, other SWI/SNF subunit proteins, and MyoD, a critical lineage determinant for skeletal muscle differentiation. We conclude that the Cn phosphatase directly impacts the expression of myogenic genes by promoting ATP-dependent chromatin remodeling and formation of transcription-competent promoters.

## INTRODUCTION

Myoblast differentiation is an essential process during skeletal muscle development where mononuclear myoblasts withdraw from the cell cycle and undergo fusion and other morphological changes to form multi-nucleated myotubes. This process is coordinated by the family of myogenic regulatory factors (MRFs) that include MyoD, myogenin, Myf5, and Mrf4 in cooperation with the MEF family of transcription factors and other auxiliary transcriptional regulators. MRFs regulate the commitment, determination, and differentiation of skeletal muscle progenitor cells. The ability of MRFs to drive the myogenic gene expression needed for differentiate requires remodeling of chromatin at target genes that depends on the recruitment of histone modifying and chromatin remodeling complexes that alter nucleosome structure and the local chromatin environment (1-3).

The SWI/SNF (<u>SWI</u>tch/<u>S</u>ucrose <u>N</u>on-<u>F</u>ermentable) complexes are large, multiprotein, ATP-dependent chromatin remodeling enzymes (4-6) that alter nucleosome structure to promote transcription, replication, recombination and repair (7-10). The chromatin remodeling activity of the SWI/SNF enzyme is required for the initiation of many developmental and differentiation programs (11-14) including activation of myogenic genes upon differentiation signaling (15, 16). Mammalian SWI/SNF complexes contain one of two related ATPase subunits, either Brahma related gene 1 (Brg1) or the ATPase Brahma (Brm), and a collection of at least 9 to 12 associated protein known as Brg1/Brm - associated factors (Bafs) (4, 17, 18). Mammalian SWI/SNF enzyme function can be influenced by the assembly of different combination of Baf subunits around the different ATPases (12, 19). Furthermore, signal transduction pathways promote specific posttranslational modifications of SWI/SNF subunit proteins that influence enzyme activity (15, 20-23). In skeletal muscle differentiation, the p38 mitogen–activated protein kinase (MAPK) phosphorylates the Baf60c subunit, which then allows the recruitment of the rest of SWI/SNF remodeling complex to myogenic promoters (24). Our group previously showed that casein kinase 2 (CK2) phosphorylates Brg1 to regulate *Pax7* expression and to promote myoblast survival and proliferation (21), while protein kinase C β1 (PKCβ1) phosphorylates Brg1, which represses chromatin remodeling function and, consequently, myogenesis (20).

Calcineurin (Cn) is a serine/threonine phosphatase that is regulated by changes in the intracellular concentration of Ca^2+^ (25). Cn is a heterodimer formed by association of catalytic subunit A (CnA) and regulatory subunit B (CnB) (26, 27). Its mechanism of action has been characterized extensively in lymphocytes, where activated Cn dephosphorylates <u>N</u>uclear <u>F</u>actor of <u>A</u>ctivated <u>T</u>-cell (NFAT) transcription factors. Dephosphorylated NFAT translocates to the nucleus and binds to promoter regions of target genes to regulate gene expression (28-31). In skeletal muscle, Cn-dependent binding of NFAT to target promoters controls skeletal muscle fiber type and primary muscle fiber number during development (32, 33) and growth of multinucleated muscle cells (34-36). Cn is also required for the initiation of skeletal muscle differentiation by mechanisms that are independent of NFAT (37, 38). More recently, we reported a novel function for Cn in chromatin remodeling. We showed that Cn is bound to Brg1 at the myogenin promoter, where it dephosphorylates Brg1 shortly after cells start the differentiation process to positively promote differentiation (20).

The aim of the current study was to explore the global effect of Cn on gene expression in myoblasts. We demonstrate that inhibition of Cn in myoblasts globally down-regulates expression of genes important for muscle structure and function. We identified four major temporal patterns of Cn-dependent gene expression. Mechanistically, we show that Cn acts as a chromatin binding regulatory protein, interacting with Brg1 to facilitate SWI/SNF enzyme and MyoD binding to myogenic gene regulatory sequences.

## MATERIALS AND METHODS

### Antibodies

Rabbit antisera to Brg1 and MyoD were previously described (39, 40). Pan-Calcineurin A (#2614), Baf170 (#12769), and Baf250A (#12354) antibodies were from Cell Signaling Technologies (Danvers, MA). Brg1 antibody (G-7, #sc-17796) was from Santa Cruz Biotechnologies (Santa Cruz, CA) and was used for western blotting and immunoprecipitation experiments.

### Cell culture

C2C12 cells were purchased from ATCC (Manassas, VA) and maintained at subconfluent densities in Dulbecco’s modified Eagle’s medium (DMEM) supplemented with 10% FBS and 1% penicillin/streptomycin in a humidified incubator at 37°C in 5% CO_2_. For differentiation, cells at > 80% confluency were switched to DMEM medium supplemented with 2% horse serum and 2 μg/ml of bovine insulin (Sigma-Aldrich, St. Louis, MO). FK506 (Cayman Chemical, Ann Arbor, MI) was added to the culture 24h before initiating differentiation and maintained in the differentiation media at 2μM. Media containing FK506 was changed every day.

Mouse satellite cells were isolated from leg muscles of 3-to 6-week old Brg1 conditional mice using Percoll sedimentation followed by differential plating as described previously (20). Mice were housed in the animal care facility at the University of Massachusetts Medical School and used in accordance with a protocol approved by the Institutional Animal Care and Use Committee. Brg1 depleted primary myoblasts expressing wild type (WT) Brg1 or Brg1 mutated at sites of PKCβ1/Cn activity were generated as described (20). Primary myoblasts were grown and differentiated as described (20) on plates coated overnight in 0.02% collagen (Advanced BioMatrix, San Diego, CA).

### RNA isolation and gene expression analysis

RNA was extracted using TRIzol Reagent (Invitrogen, Carlsbad, CA) and the yield determined by measuring OD_260_. 1µg of total RNA was subjected to reverse transcription with a QuantiTect Reverse Transcription Kit (Qiagen, Germantown, MD). The resulting cDNA was used for quantitative PCR using a Fast SYBR green master mix (Applied Biosystems, Foster City, CA). Amplification reactions were performed in duplicate in 10 μl final volume that included the following: 25 ng of template, 0.3 μM primers, 2× SYBR Green Master Mix. Reactions were processed in QuantStudio3 (Applied Biosystems). Δ*C*_t_ for each gene was calculated and represents the difference between the *C*_t_ value for the gene of interest and that of the reference gene, *Eef1A1*. Fold-changes were calculated using the 2^−ΔΔ*C*_t_^ method (41). Primer sequences and their accession numbers of PCR products are shown in **Supplementary Table S1**.

### RNA-Seq and Data Analysis

Total RNA from C2C12 cells treated with FK506 or with DMSO was isolated from proliferating and differentiating cultures (time 0, 24h and 72h) with TRIzol, and libraries were constructed as described (42). The libraries were sequenced using the Illumina HiSeq 1500 and the resulting reads were mapped onto the reference mouse genome (GRCm38) by HISAT2 (ver. 2.2.6) (43). Read counting per gene was performed with HTseq (ver. 0.6.1) (44) such that duplicates in unique molecular identifiers were discarded. After converting UMI counts to transcript counts as described (45), differentially expressed genes, (those with adjusted p-value <0.1) were extracted by the R library DESeq2 (version 1.10.1) (46). The differentially expressed genes and cluster analyses are listed in **Supplementary Tables S2 and S3**. Gene ontology (GO) term identification was performed on metascape: http://metascape.org. Cluster gene analysis was performed using clusterProfiler (ver. 3/10.0) software (47). RNA-seq data were deposited at the Gene Expression Omnibus (GEO) database under accession number GSE125914. The reviewer access token is krexscswllodxwn.

### Chromatin immunoprecipitation (ChIP) assay

Chromatin immunoprecipitation assays were performed as previously described (20, 48) with some modifications. Briefly, cells (4 × 10^6^) were cross-linked with 1% formaldehyde (Ted Pella Inc., Redding, CA) for 10 minutes at room temperature. After quenching the formaldehyde with 125 mM glycine for 5 minutes, fixed cells were washed twice with PBS supplemented with protease inhibitor cocktail and lysed with 1 ml buffer A (10 mM Tris HCl, pH 7.5, 10 mM NaCl, 0.5% NP40, 0.5 μM DTT and protease inhibitors) by incubation on ice for 10 minutes. The nuclei were pelleted, washed with 1 ml of buffer B (20 mM Tris HCl, pH 8.1, 15 mM NaCl, 60 mM KCl, 1 mM CaCl_2_, 0.5 μM DTT) and incubated for 30 minutes at 37°C in the presence of 1000 gel units of MNase (#M0247S, NEB, Ipswich, MA) in 300 μl volume of buffer B. The reaction was stopped by adding 15 μl of 0.5 M EDTA. Nuclei were pelleted and resuspended in 300 μl of ChIP buffer (100 mM Tris HCl, pH 8.1, 20 mM EDTA, 200 mM NaCl, 0.2 % sodium deoxycholate, 2% Triton X 100 and protease inhibitors), sonicated for 10 minutes (30 sec-on/ 30 sec-off) in a Bioruptor UCD-200 (Diagenode, Denville, NJ) and centrifuged at 15,000 rpm for 5 minutes. The fragmented chromatin was between 200–500 bp as analyzed on agarose gels. Chromatin concentration was measured using Qubit 3 (Invitrogen). After preclearing with protein A agarose, chromatin (2-4 µg) was subjected to immunoprecipitation with specific antibodies listed above, or with anti-IgG as negative control at 4°C overnight, and immunocomplexes were recovered by incubation with protein A agarose magnetic beads (Invitrogen). Sequential washes of 5 minutes each were performed with buffers A-D (Buffer A: 50 mM Tris pH 8.1, 10 mM EDTA, 100 mM NaCl, 1% Triton X 100, 0.1% sodium deoxycholate; Buffer B: 50 mM Tris pH 8.1, 2 mM EDTA, 500 NaCl, 1% Triton-X100, 0.1% sodium deoxycholate; Buffer C: 10 mM Tris pH 8.1, 1 mM EDTA, 0.25 M LiCl_2_; 1% NP-40, 1% sodium deoxycholate; Buffer D: 10 mM Tris pH 8.1, 1 mM EDTA), immune complexes were eluted in 100 µl of elution buffer (0.1 M NaHCO_3_, 1% SDS) for 30 minutes at 65°C, incubated with 1 μl of RNnase A (0.5 mg/ml) for 30 minutes at 37°C, and reverse cross-linked by adding 6 μl of 5M NaCl and 1 μl of proteinase K (1 mg/ml) overnight at 65°C. DNA was purified using the ChIP DNA clean & concentrator kit (Zymo Research, Irvine, CA). Bound DNA fragments were analyzed by quantitative PCR using the SYBR Green Master Mix. Quantification was performed using the fold enrichment method (2^−(Ct sample − Ct IgG)^) and shown as relative to a control region, the promoter for the *Pdx1* gene. Primer sequences are listed in **Supplemental Table S1**. Primer positions for each promoter are shown schematically in **Supplemental Figure S1**.

### Immunoprecipitation

100 mm dishes of 24h differentiated C2C12 cells treated with FK506 or with DMSO or 100 mm dishes of primary myoblasts expressing wildtype or Brg1 mutants were washed with ice-cold PBS twice and lysed in 0.5 ml of lysis buffer (50 mM Tris-HCl, pH 7.5, 150 mM NaCl, 1% Nonidet P-40, 0.5% sodium deoxycholate, 1mM CaCl_2_ and supplemented with complete protease inhibitors). Lysates were cleared by centrifugation and precleared with PureProteome Protein A magnetic beads (Millipore) for 2 hours at 4°C. Next, cell extract was incubated with 10 μl of Pan-Calcineurin A antibody (Cell Signaling Technologies) or 10 μl of Brg1 antisera overnight at 4°C, followed by incubation with 100 μl with PureProteome Protein A mix magnetic beads. After extensive washing of beads with washing buffer (24 mM Tris-HCl, pH 7.5, 300 NaCl, 0.5% NP-40 and 1mM CaCl_2_), precipitated proteins were eluted in Laemmli buffer and detected by western blot analysis using chemiluminescent detection.

### Statistical analysis

All quantitative data are shown as mean +/- the standard deviation (SD) of at least three (n=3) biological replicates for each experiment. Statistical analysis was performed using GraphPad Prism Student’s t-test (GraphPad Software Inc.). For all analyses, a *P*-value of less than 0.05 was considered to be statistically significant.

## RESULTS

### RNA-seq identification of genes differentially expressed in myoblasts treated with calcineurin inhibitor FK506

To better understand the involvement of calcineurin in the gene regulation program underlying myogenesis, we compared the transcriptomes of C2C12 myoblasts at three time points of differentiation (0, 24, and 72 h post-induction of differentiation) in cells treated with the Cn inhibitor, FK506, with those of control cells. FK506 inhibits the phosphatase activity of Cn by binding to the immunophilin FKB12; the drug-bound FKB12 binds to and blocks Cn function (28). Myogenic gene expression is temporally regulated, with different genes expressed with different kinetics during differentiation (49-51). We analyzed gene expression 24 h and 72 h after the induction of differentiation to distinguish the impact of Cn inhibition on genes expressed at early or late times of myogenesis.

At 24h post-differentiation, 156 genes were differentially expressed in the presence of FK506 [false discovery rate (FRD) < 0.05]. Fewer genes were up-regulated (53) than were downregulated (103). At the later time point of differentiation (72h), 648 genes were differentially expressed. More genes were up-regulated (373) than were downregulated (275) (**Fig. 1A, B**). In confluent myoblasts at the onset of differentiation (0h), only 21 genes were differentially expressed in the presence of FK506. The complete RNA-seq analysis of differentially expressed genes is shown in **Supplementary Table S2**. Gene ontology analysis of genes downregulated by calcineurin inhibition at the 24 and 72h time points demonstrated that these genes are involved in muscle differentiation and related functions (**Fig. 1C** and **Supplementary Fig. S2**). Specific myogenic genes were identified and labeled on the volcano plots in **Fig. 1A.** In contrast, we observed up-regulation of genes associated with ossification, response to interferon and viruses, and cardiovascular and blood vessel development. These results are potentially consistent with documented roles for Cn in the regulation of NFAT function in these processes (52-54). We performed *de novo* motif analysis on the differentially expressed genes. The most significantly enriched motifs at the 24h post-differentiation were for Mef2 transcription factors and E-boxes bound by the MRFs Myf5, myogenin and MyoD (**Fig. 1D**). These findings indicate that genes that require calcineurin for expression are also regulated by myogenic regulatory factors, strongly reinforcing the connection between Cn and expression of the myogenic gene program. At the 72h time point, the most significantly enriched motif was the interferon regulatory factor (IRF) binding sequence **(Fig. 1D)**. This is consistent with the identification of G0 terms including response to interferons and with findings that IRFs regulate expression of vascular cell adhesion molecule 1 (VCAM-1) receptor that mediates cell-cell adhesion and is important for myoblast fusion (55, 56). We also note the continued presence of MRF and Mef2 protein binding sites at Cn-regulated genes at late times of differentiation.

**Figure 1:**
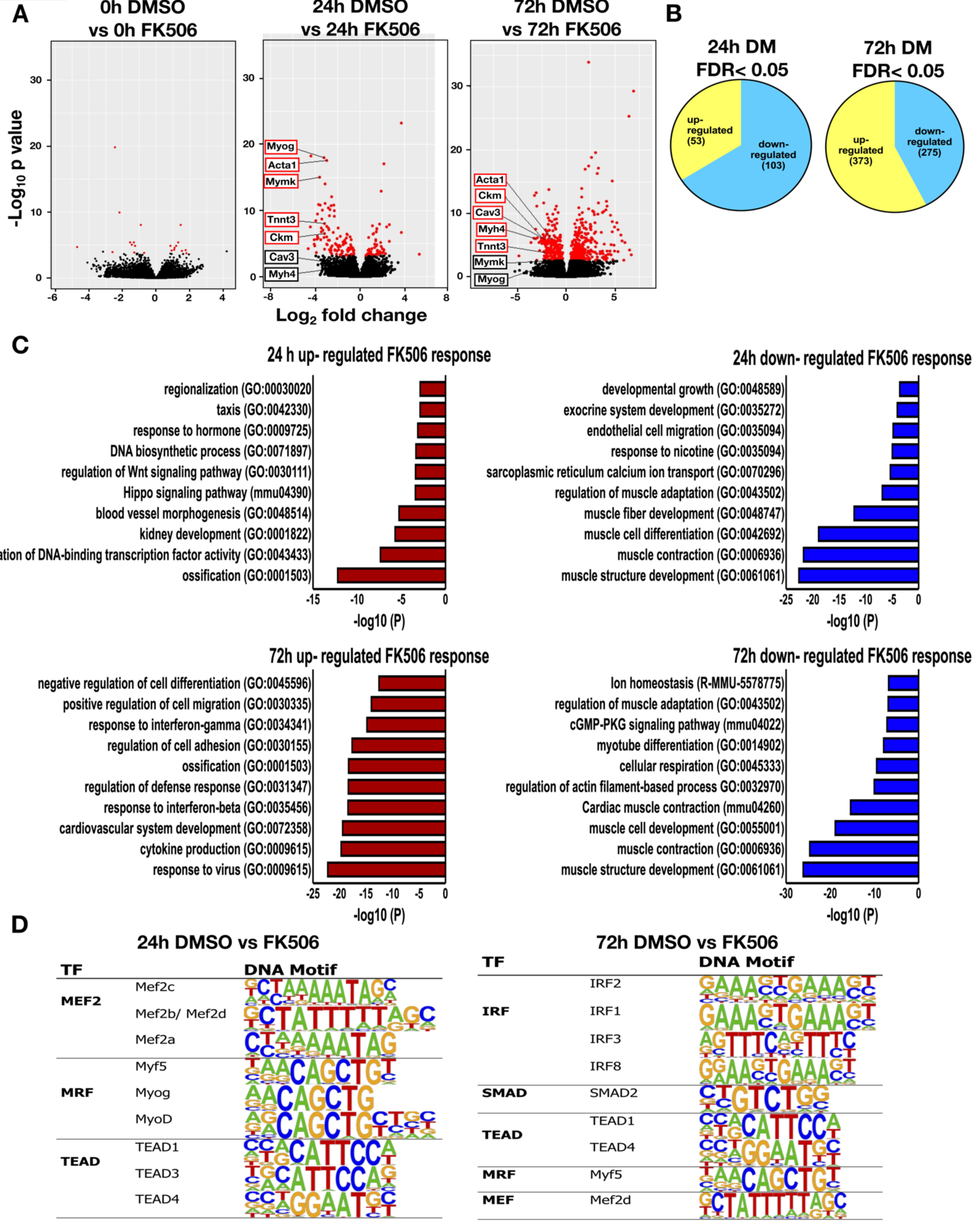
Differential gene expression in myoblasts treated with Cn inhibitor. (A) Volcano plots displaying differentially expressed genes between control (DMSO) treated and Cn inhibitor (FK506) treated differentiated C2C12 cells. The y–axis corresponds to the mean expression value of log 10 (p-value), and the x-axis displays the log2 fold change value. The red dots represent the up-and down-regulated transcripts between DMSO-and FK506-treated samples (False Discovery Rate (FDR)<0.05). The black dots represent the expression of transcripts that did not reach statistical significance (FDR>0.05). **(B)** A Venn diagram displaying the number of genes up-and down-regulated by FK506 treatment at 24 and 72h post-differentiation. **(C)** Gene ontology analyses on genes differentially expressed by FK506 treatment for 24 and 72h post-differentiation. **(D)** Transcription factor binding motifs identified within 1kb upstream of the TSS of genes differentially expressed by FK506 treatment in cells differentiated for 24 and 72h.

### Global expression analysis reveals four major temporal gene expression patterns that are dependent on Cn

We continued our analysis by identifying the groups of genes that showed treatment–specific changes in gene expression over time. In total, 308 genes were identified. We analyzed the expression patterns of these differentially expressed genes using Cluster Profiler (47). Genes differentially expressed over time were clustered into 4 major groups, graphically presented by heat map (**Fig. 2A**) and expression kinetics (**Fig. 2B**). We performed GO analysis for the differentially expressed genes in these 4 clusters to gain additional insight into the biological processes regulated by calcineurin during myoblast differentiation (**Fig. 2C**). Cluster 1 included 95 genes that were down-regulated during differentiation but were unchanged in the presence of the Cn inhibitor. GO terms significantly enriched for this group of differentially expressed genes related to cell migration, motility and adhesion. In addition, Cluster 1 included Inhibitor of differentiation (Id) proteins 1, 2 and 3 (**Supplementary Table S3**). Id proteins interact with MyoD and related MRFs prior to differentiation to repress transcription activation activity. Id1 and Id3 have previously been identified as genes repressed by Cn during the activation of skeletal muscle cell differentiation (38). Cluster 2 contained 80 genes that were up-regulated during myoblast differentiation but were inhibited or delayed in the presence of FK506. GO terms significantly enriched for Cluster 2 genes were related to muscle structure and function. Cluster 3 was composed of 108 genes that were expressed at relatively constant levels across the differentiation time course but that were significantly up-regulated in the presence of FK506 expression. GO analysis of Cluster 3 genes identified genes implicated in immune response of cells. Cluster 4 had 25 genes with an unusual profile. These genes were up-regulated in the presence of FK506 in proliferating cells and in cells at the onset of differentiation, but were down-regulated by FK506 treatment as differentiation advanced. No specific GO terms were identified for this Cluster 4. A complete list of genes in each of the clusters with their log fold change and FDR values are shown in **Supplementary Table S3**. These data suggests that calcineurin regulates multiple and diverse groups of genes during myoblast differentiation.

**Figure 2:**
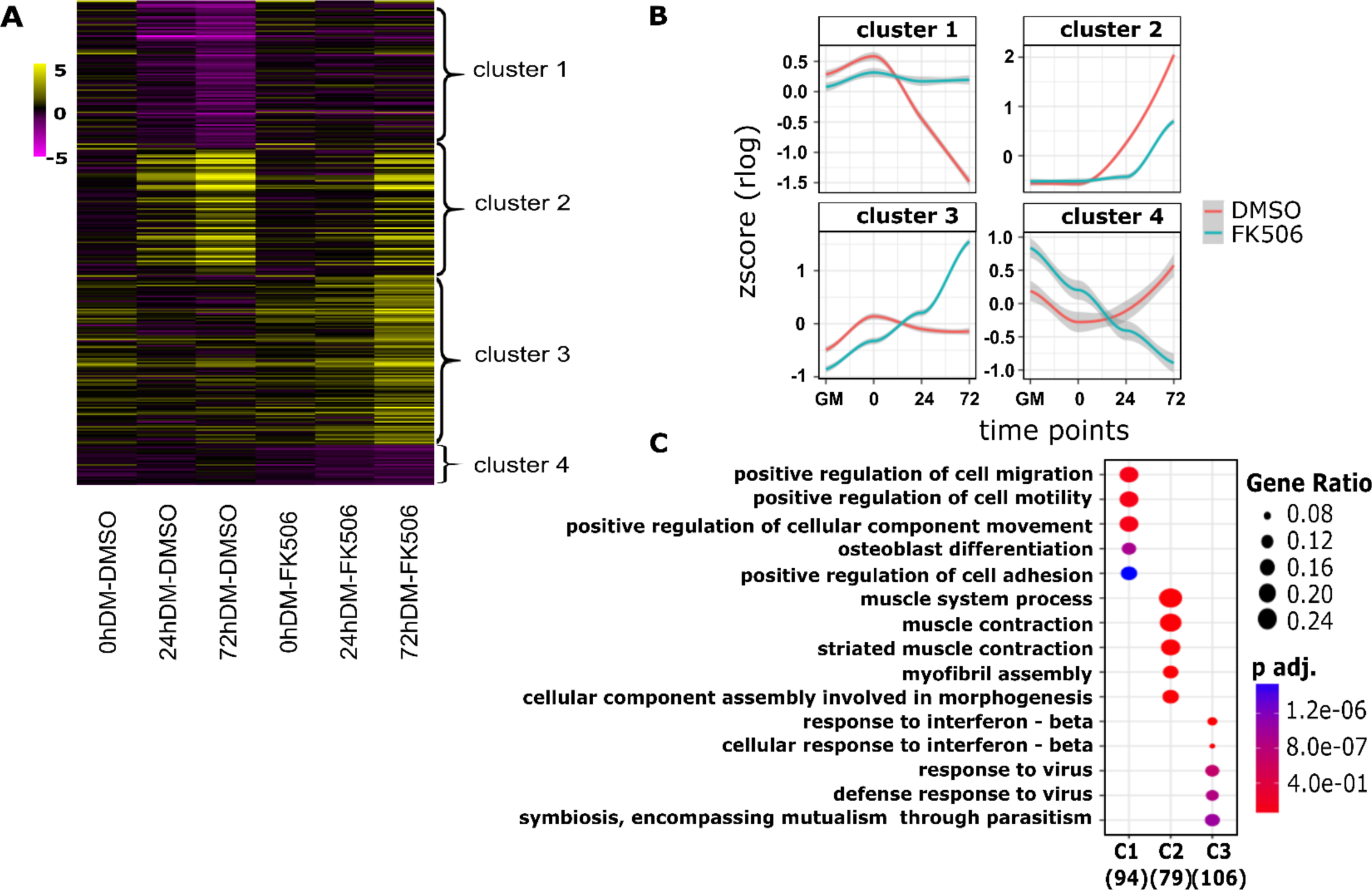
Cluster analysis of differentially expressed genes at three differentiation time points in myoblasts treated with Cn inhibitor. **(A)** The heat map comparing differential expression of 308 FK506 treatment-specific genes, categorized in four different clusters. Each column represents an experimental sample (times 0, 24 and 72h in differentiation medium (DM)) compared to the proliferating myoblast sample cultured in growth media (GM). Each row represents a specific gene. The colors range from yellow (high expression) to magenta (low expression) and represent the relative expression level value log2 ratios. **(B)** Kinetic expression patterns of the four clusters of genes. **(C)** Gene ontology analysis of differentially expressed genes within clusters 1-3 (C1, C2, C3) identified the top enriched GO terms with the corresponding enrichment *p* values and gene ratio.

For further analysis, we focused on genes that were down-regulated in the presence of FK506 and are important for muscle structure and/or function (**Fig. 1A**). We first measured mRNA expression level of several such genes to validate RNA-seq results; *Myog, Ckm, Myh4, Mymk, Tnnt3, Acta1 and Cav3* were analyzed by RT-qPCR. As expected, each of these genes were increased over the course of the differentiation time course and each was significantly down-regulated by exposure to the Cn inhibitor, FK506 (**Fig. 3**). These results confirmed that inhibition of Cn during myoblast differentiation impairs expression of multiple myogenic genes that are normally activated during differentiation.

**Figure 3:**
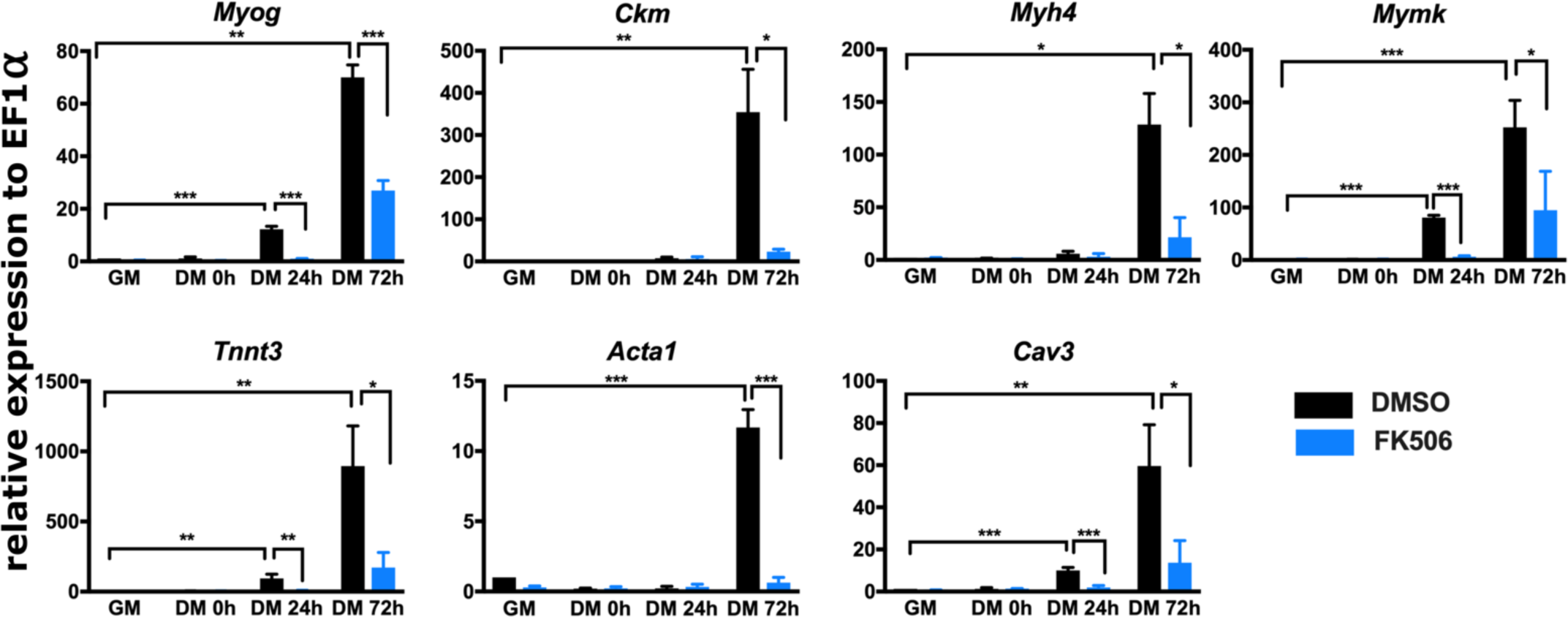
Cn regulates the expression of myogenic genes during myoblast differentiation. Real-time RT-PCR showed that expression of *Myog, Ckm, Myh4, Mymk, Tnnt3, Acta 1*, and *Cav 3* is down-regulated in FK506-treated C2C12 cells. GM, proliferating cells in growth medium. DM, differentiation medium for the indicated time in hours (h). Data are the average of three or more independent samples performed in duplicate and are presented as the mean +/- SD. Expression in DMSO-treated GM samples were set to 1 and other values are relative to that sample. * *p* ≤ 0.05, ** p ≤ 0.001, ***p≤ 0.0001 vs. GM or vehicle by Student’s t-test.

### Inhibition of calcineurin function prevents its binding to myogenic promoters

We previously showed that Cn associates with Brg1 and binds to the myogenin promoter early after the start of the differentiation process (20). We hypothesized that the same model might be true for other myogenic genes. Alternatively, Cn may directly regulate the myogenin promoter but indirectly regulate other downstream genes and via the dependency of other genes on myogenin for completing the differentiation process (57, 58). We performed chromatin immunoprecipitation assays on the same panel of myogenic genes examined in Fig. 3. PCR primers for ChIP were designed to amplify the E-box containing regions within 1.5 Kb upstream of the TSS (**Supplementary Figure S1**). Cn did not interact with myogenic genes regulatory sequences in proliferating myoblasts (GM), was weakly bound in undifferentiated cells at the start of differentiation (0h DM), and was bound at all myogenic gene promoters in differentiated (48h DM) cells (**Fig. 4**). We conclude that Cn binding to myogenic promoters is a general occurrence and FK506-mediated inhibition of Cn function impairs its ability to interact with target gene promoters. The results suggest that Cn is a direct regulator of many, if not all, myogenic genes.

**Figure 4:**
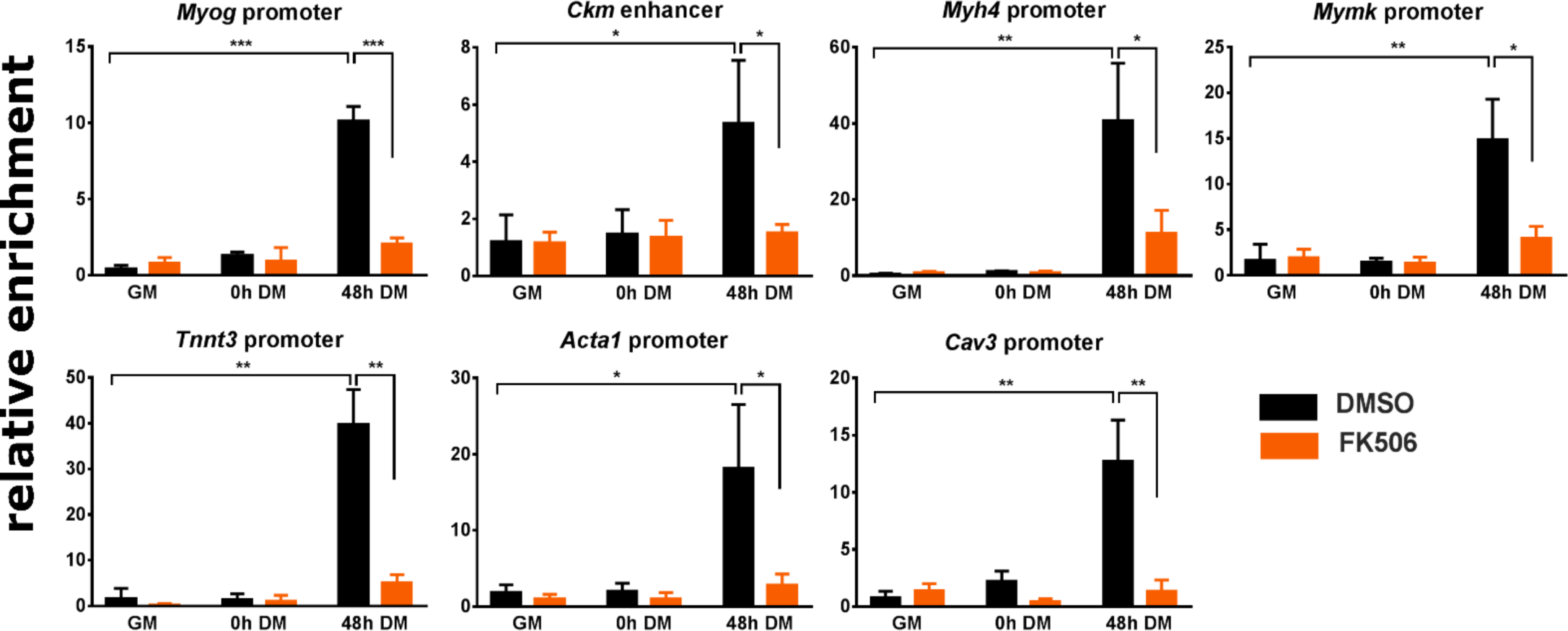
Cn binds to E-box containing regulatory sequences of myogenic genes during myoblast differentiation. Cn binding is reduced by FK506 treatment. ChIP assays were performed for Cn binding in C2C12 cells. GM, proliferating cells in growth medium. DM, differentiation medium for the indicated time in hours (h). Relative enrichment was defined as the ratio of amplification of the PCR product normalized to control IgG and is shown relative to amplification of a non-specific control promoter region. The data are average of at least 3 independent experiments performed in triplicate +/- SD. * p ≤ 0.05, ** p ≤ 0.01, ***p≤ 0.001 vs. GM or vehicle by Student’s t-test.

### Cn inhibition blocks the interaction of the ATPase Brg1 and other subunits of the mammalian SWI/SNF complex with myogenic promoters

We also performed ChIP assays to determine whether Brg1 recruitment to myogenic regulatory sequences was dependent on calcineurin. We observed recruitment of Brg1 to regulatory sequences in differentiated (48h DM) cells at all the tested myogenic genes. In cells treated with FK506, Brg1 binding was significantly diminished at all promoters (**Fig. 5A**). A subset of the myogenic promoters was tested for the binding of Baf170, and Baf250A/Arid1A, which are other subunits of the mammalian SWI/SNF enzyme complex. The results show that binding of these other subunits paralleled Brg1 binding in that Cn inhibition blocked interaction of these proteins to the promoters (**Fig. 5B)**. These findings indicate that Cn activity is necessary for the interaction of mammalian SWI/SNF chromatin remodeling complexes with regulatory sequences of myogenic genes.

**Figure 5:**
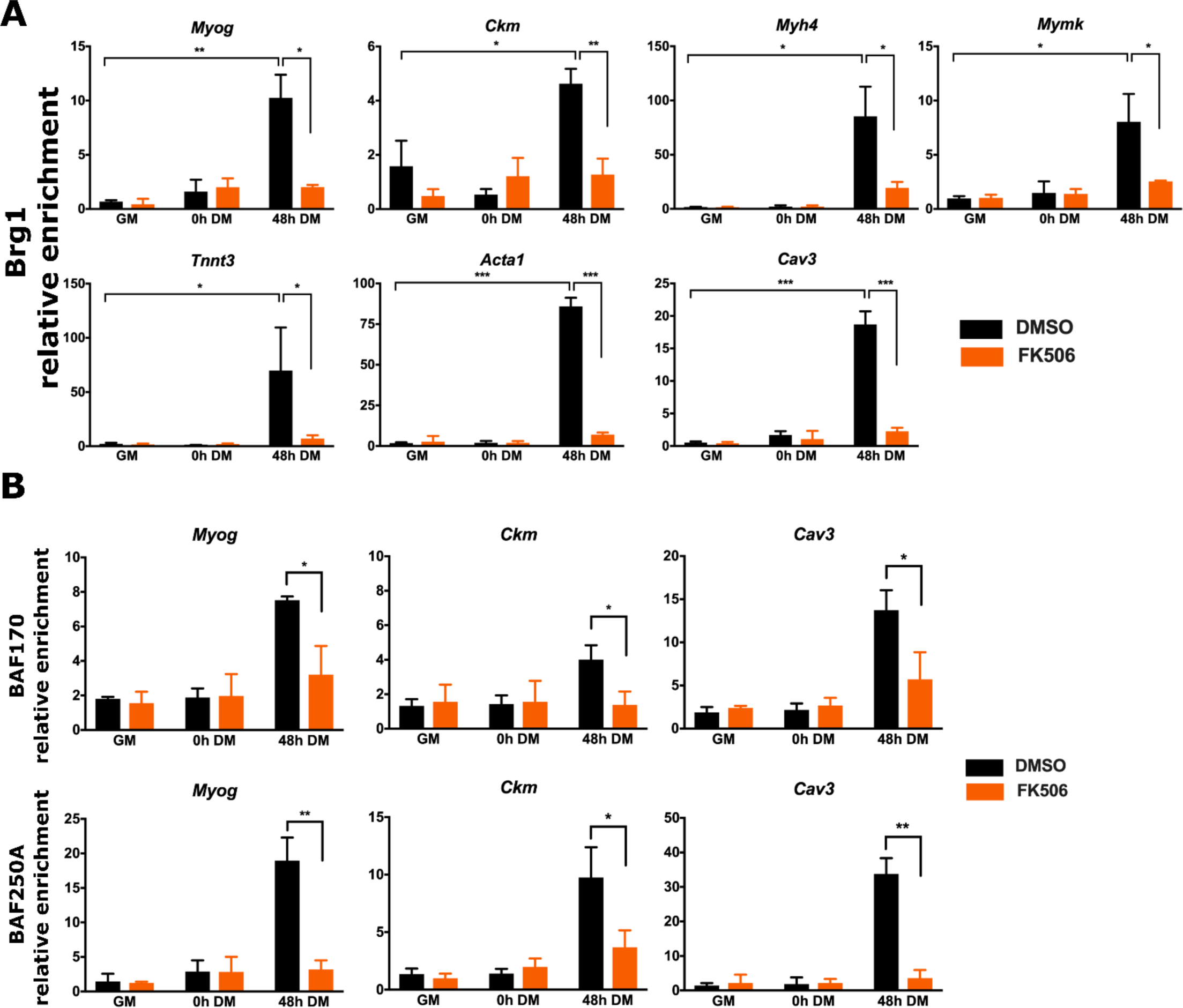
Cn inhibition reduces binding of the SWI/SNF subunits Brg1 (A), Baf170 and Baf250A (B) to E-box containing regulatory sequences of myogenic genes during myoblast differentiation. ChIP assays were performed for Brg1, Baf170 and Baf250A binding in C2C12 cells. GM, proliferating cells in growth medium. DM, differentiation medium for the indicated time in hours (h). Relative enrichment was defined as the ratio of amplification of the PCR product normalized to control IgG and is shown relative to amplification of a non-specific control promoter region. The data are average of at least 3 independent experiments performed in triplicate +/- SD. * p ≤ 0.05, ** p ≤ 0.01, ***p≤ 0.001 vs. GM or vehicle by Student’s t-test.

### Mutation of Brg1 sites of Cn activity prevents its interaction with myogenic gene promoters

In previous work, we showed that PKCβ-mediated phosphorylation of Brg1 prior to the onset of differentiation was counteracted by Cn-mediated dephosphorylation of Brg1 immediately after the onset of differentiation (20). Serine amino acids targeted by PKCβ/Cn mapped to 5’ and 3’ of the Brg1 bromodomain. Mutation of these sites to the phosphomimetic amino acid glutamate (SE) prevented myogenesis, whereas mutation to the non-phosphorylatable amino acid alanine (SA) had no effect on differentiation (20). These experiments used primary myoblasts derived from Brg1-deficient mice that were reconstituted with wildtype (WT)-, SA-, or SE-Brg1. We performed ChIP experiments in differentiating cells and showed that Cn and Brg1 are bound to myogenic promoters in myoblasts expressing WT-Brg1 (**Fig. 6A, B**). The SE-Brg1 mutant was incapable of binding; the repressive phosphorylation of Brg1 caused by PKCβ is mimicked by the glutamate substitutions, rendering Cn incapable of activating Brg1. As expected, the SA-Brg1 mutant, which cannot be phosphorylated at the relevant PKCβ target amino acids, bound Brg1 and Cn to myogenic regulatory sequences (**Fig. 6A, B**). These results are consistent with those obtained with the Cn inhibitor, and they reinforce the conclusion that Cn function regulates Brg1 binding to chromatin at myogenic gene regulatory sequences.

**Figure 6:**
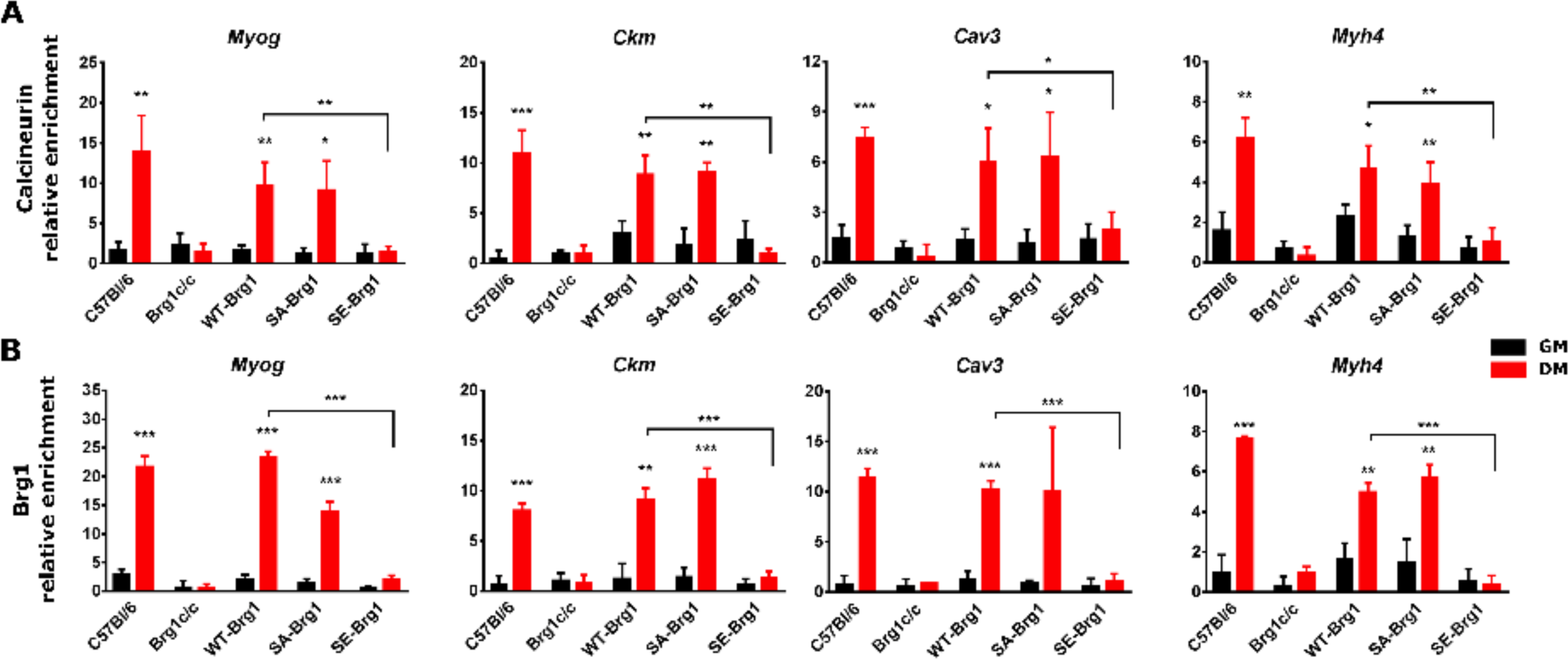
Phosphomimetic mutation of Brg1 amino acids that are dephosphorylated by Cn reduces Cn and Brg1 binding to myogenic promoters in differentiating myoblasts. ChIP assays were performed for Cn or Brg1 binding in primary mouse myoblasts (C57Bl/6), in primary mouse myoblasts deleted for the gene encoding Brg1 (Brg1c/c), or in primary mouse myoblasts deleted for the gene encoding Brg1 that are expressing a wildtype (WT-Brg1), Brg1 containing a non-phosphorylatable amino acid at Cn-targeted sites (SA-Brg1), or Brg1 containing a phosphomimetic amino acid at Cn-targeted sites (SE-Brg1). Samples were collected from proliferating cells in growth medium (GM) or at 24h post-differentiation (DM). Relative enrichment was defined as the ratio of amplification of the PCR product normalized to control IgG and is shown relative to amplification of a non-specific control promoter region. The data are average of 3 independent experiments performed in triplicate +/- SD. * p ≤ 0.05, ** p ≤ 0.01, ***p≤ 0.001 by Student’s t-test.

### Cn inhibition does not impact the ability of Cn and Brg1 to interact

We previously showed that Cn and Brg1 can be co-immunoprecipitated from cell lysate of differentiating cells (20). Inhibition of Cn function with FK506 had no impact on the ability of these proteins to be isolated in complex with each other (**Fig. 7A**). As a complement to this experiment, we looked at the interaction of Cn with Brg1 in differentiating myoblasts expressing WT-, SA-, or SE-Brg1. Cn could interact with WT-, SA-, and SE-Brg1 mutants (**Fig. 7B**), despite the observation that SE-Brg1 was not competent for interaction with chromatin. These results indicate that inhibition of Cn function and mutation of the sites of Cn activity on Brg1 do not affect the interaction that exists between these regulatory proteins. The continued existence of Brg1 protein in the presence of the Cn inhibitor and when Cn-targeted residues are mutated to alanine or glutamine suggests that the lack of appropriate phosphorylation or dephosphorylation does not have significant impact on the steady-state levels of Brg1 and therefore is unlikely to be a major regulator of protein stability.

**Figure 7:**
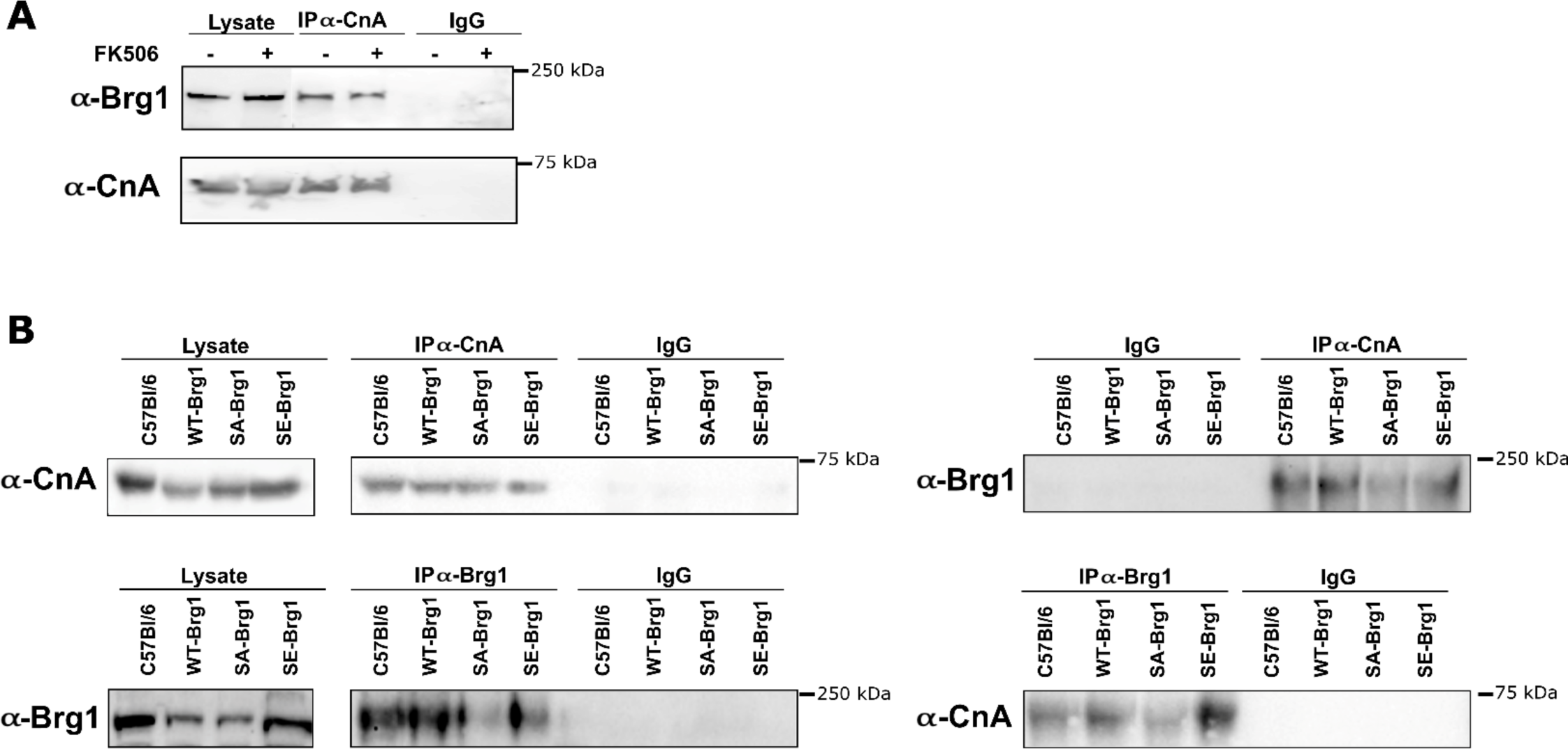
Interaction between Cn and Brg1 is not affected by Cn inhibition (A) or mutation of Brg1 amino acids that are targeted by Cn (B). **(A)** Co-immunoprecipitation of Cn and Brg1 from cell lysates from differentiated C2C12 cells treated with FK506. **(B)** Co-immunoprecipitation of Cn and Brg1 from cell lysates from 24h differentiated primary C57Bl/6 myoblasts and from primary mouse myoblasts deleted for the gene encoding Brg1 that are expressing a wildtype (WT-Brg1), Brg1 containing non-phosphorylatable amino acids at Cn-targeted sites (SA-Brg1), or Brg1 containing phosphomimetic amino acids at Cn-targeted sites (SE-Brg1). Cell lysate from each IP (2.5% of input) served as a loading controls. The experiments were performed 3 times and representative gels are shown.

### Inhibition of calcineurin blocks MyoD binding to regulatory sequences of myogenic genes

Recruitment of MyoD to myogenic promoters prior to the onset of differentiation can be accomplished by different mechanisms, including gene-specific mechanisms (24, 40, 59). The continued presence of MyoD on myogenic promoters after the onset of differentiation requires the Brg1 ATPase (15, 40). We would therefore predict that the inhibition of calcineurin would affect the interaction of MyoD with myogenic gene regulatory sequences. We assessed MyoD enrichment at the regulatory sequences of tested genes by ChIP. As shown in **Fig. 8**, we observed enhanced enrichment of MyoD at all analyzed gene promoters in differentiated cells compared to enrichment prior to or at the onset of differentiation. Recruitment of MyoD at these regulatory sequences was attenuated by the calcineurin inhibitor. These results support the conclusion that calcineurin is necessary for the stable binding of MyoD to the myogenic gene regulatory sequences during differentiation.

**Figure 8:**
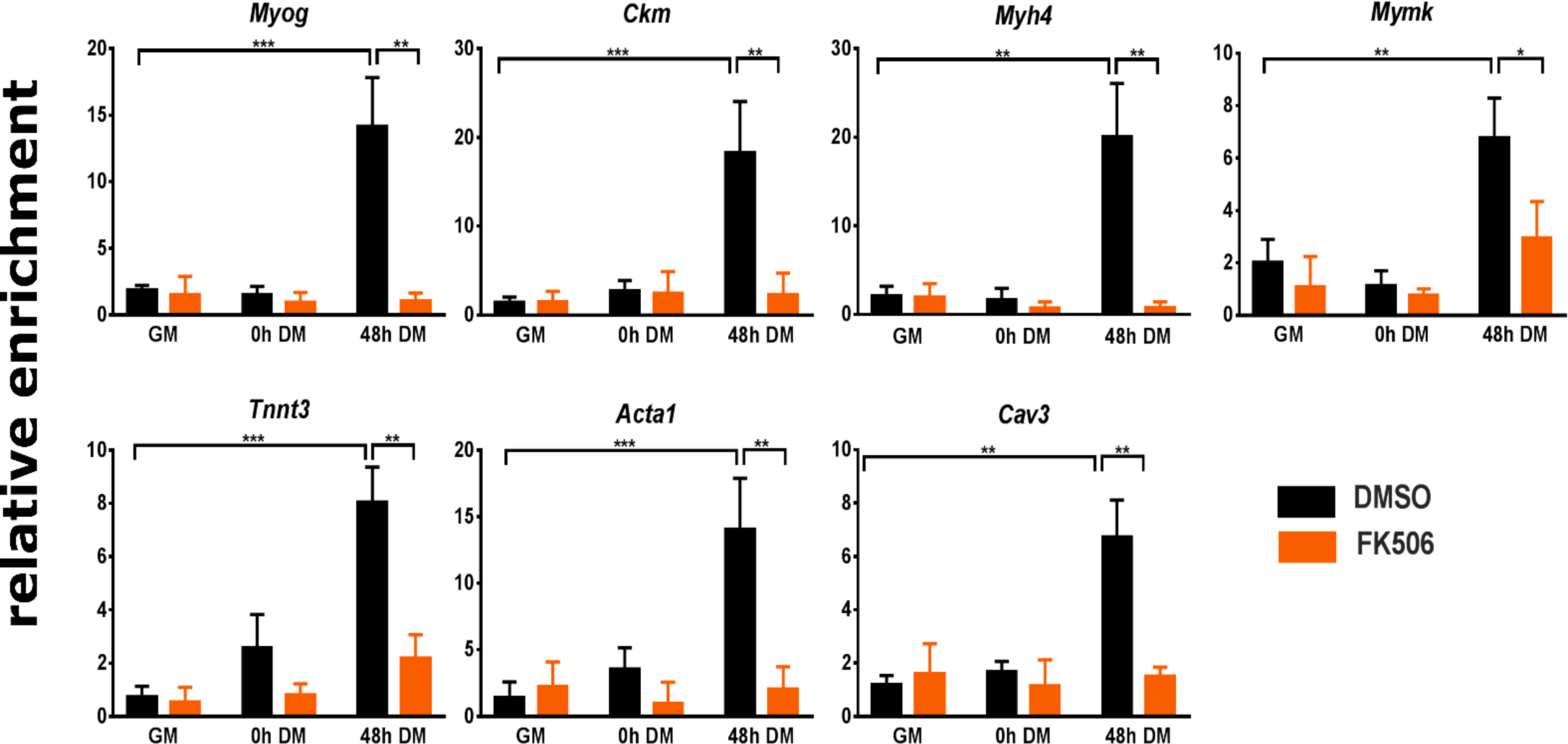
Inhibition of Cn reduced MyoD binding to regulatory sequences of myogenic genes during myoblast differentiation. ChIP assays were performed for MyoD binding in C2C12 cells. GM, proliferating cells in growth medium. DM, differentiation medium for the indicated time in hours (h). Relative enrichment was defined as the ratio of amplification of the PCR product normalized to control IgG and is shown relative to amplification of a non-specific control promoter region. The data are average of at least 3 independent experiments performed in triplicate +/- SD. * p ≤ 0.05; ** p ≤ 0.01, *** p≤ 0.001 vs. GM or vehicle by Student’s t-test.

## DISCUSSION

### Cn broadly contributes to the activation of the myogenic gene expression program during differentiation

The data presented here demonstrate that Cn plays a general role in regulating myogenic gene expression during the myoblast differentiation. Its mechanism of action is via direct promoter binding and dephosphorylation of the Brg1 ATPase of the mammalian SWI/SNF chromatin remodeling enzyme, which regulates the ability of Brg1 and other SWI/SNF enzyme subunits to stably associated with myogenic promoters during differentiation. Failure of this regulatory step prevents the required enzymatic remodeling of promoter chromatin structure and subsequent gene activation during differentiation.

The consistent observation of Cn binding to each of the myogenic promoters assayed suggests that Cn is directly required for the activation of each target gene. The alternative hypothesis was that Cn required indirectly via a requirement for activation of myogenin, which is required for activation of myogenic gene products that promote terminal differentiation (60, 61). In prior work examining other cofactors of myogenic gene expression, we determined that the SWI/SNF chromatin remodeling enzyme was required for the expression of both myogenin and subsequent gene expression (57). These new results spatially link Cn and SWI/SNF enzyme binding to myogenic promoters, which is consistent with Cn function being required for SWI/SNF enzyme function. In contrast, the Prmt5 arginine methyltransferase is required for myogenin activation, but ectopic expression of myogenin promoted myogenic gene expression and differentiation even in the absence of Prmt5 (62), indicating that the requirement for Prmt5 in later stages of the myogenic gene expression cascade was indirect.

The promoter binding capability of Cn is a novel function that is poorly understood. There is no evidence that Cn contains a recognized DNA binding domain, raising the possibility that it binds chromatin indirectly. The absence of Cn binding in the presence of the Cn inhibitor suggests the possibility of auto-dephosphorylation as a necessary pre-requisite. However, prior studies indicate that autodephosphorylation is slow and that phosphorylated Cn is more efficiently dephosphorylated by protein phosphatase IIa (63). An alternative hypothesis is that Cn inhibition alters the structure or function of a Cn binding partner, which directly or indirectly results in loss of association with chromatin.

### Regulation of SWI/SNF enzyme by subunit composition and by phosphorylation state

The diversity of mammalian SWI/SNF enzyme complex formation is due in part to the existence of several subunits that exclusively or predominantly associate with subsets of enzyme complexes (12, 19). The Baf250A subunit exists in a subset of SWI/SNF complexes known as a SWI/SNF-A or BAF, which contains several unique subunits not found in the other major subfamily of SWI/SNF complexes, referred to as SWI/SNF-B or PBAF complexes (64). The presence of Baf250A at each of the promoters assays suggests that the A/BAF complex may be the functionally relevant enzyme to promote skeletal muscle differentiation.

The literature on which specific enzyme complex(es) act at muscle-specific promoters is limited. Brg1 has been identified at many myogenic promoters. Interference with Brg1 function through expression of a dominant negative enzyme, injection of specific antibodies, or via knockdown blocks myogenic gene expression and differentiation (15, 16, 24, 40, 65), but these data do not distinguish between different types of SWI/SNF enzymes. The Brm ATPase binds to the myogenin promoter in isolated mouse myofibers, but not in isolated satellite cells (66), however, knockdown of Brm in cultured myoblasts had limited effect on differentiation-specific gene expression while instead affecting cell cycle withdrawal (65). Those data indicate that A/BAF complex function is a necessary pre-requisite for myoblast differentiation. In mouse heart development, the B/PBAF specific subunits BAF200 and BAF180 are required (67-69), but Baf250A knockout in mouse neural crest leads to embryonic death due to defective cardiac development (70). Despite the intriguing ramifications of thousands of different potential SWI/SNF enzyme compositions, comparison of complexes formed by A/BAF and B/PBAF-specific subunits in the same cell type showed that genomic binding sites and transcriptionally responsive genes largely overlapped, leading to the conclusion that the regulation of gene expression by SWI/SNF enzymes is due to the combined effect of multiple SWI/SNF enzymes (71). Muscle development and differentiation may similarly rely on multiple SWI/SNF enzyme assemblies and may not be attributable to one specific enzyme complex.

Regulation of SWI/SNF chromatin remodeling enzyme activity, via control of the phosphorylation state of different proteins within the enzyme complex, is an emerging complexity that adds to the alarming complexity posed by the thousands of potential combinatorial assemblies of the enzyme complex from its component subunit proteins (12, 19). Nevertheless, the evidence for this additional layer of regulation is clear. Amino acids 5’ and 3’ to the Brg1 bromodomain are phosphorylated by PKCβ1 in proliferating myoblasts and dephosphorylated by Cn after the onset of differentiation signaling. Failure to remove the phosphorylation prevents remodeling enzyme function and differentiation, while mutation of these amino acids to prevent phosphorylation permits function even in the presence of a Cn inhibitor (20). Here we demonstrate the generality of the requirement for Cn-mediated facilitation of Brg1 function. The PKCβ1/Cn axis is joined by p38 kinase-mediated phosphorylation of the Baf60c subunit that permits assembly of the SWI/SNF enzyme complex on myogenic promoters (24). Most of the SWI/SNF subunits have been characterized as phosphoproteins (72), suggesting that regulation of activity in response to differentiation signaling may be influenced by other kinases and phosphatases as well.

The consequences of phosphorylation and dephosphorylation of the PKCβ1/Cn-targeted amino acids on Brg1 structure remain to be determined. The fact that Brg1 protein remains present when the relevant amino acids are mutated or when exposed to Cn inhibitor indicates that the phosphorylation state of these amino acids does not significantly contribute to Brg1 protein stability. The failure of other SWI/SNF subunits to remain associated with myogenic gene chromatin suggests that the phosphorylation state may control Brg1 protein conformation, interactions with chromatin, interactions with other SWI/SNF enzyme subunits, and/or interactions with other cofactors that contribute to enzyme complex stability or chromatin binding. Additional characterization will improve our understanding of Brg1 and SWI/SNF chromatin remodeling enzyme function in differentiation and may also inform studies on the role of Brg1 in oncogenesis, where Brg1 can be either mutated or overexpressed without mutation in different types of cancer (73, 74).

## ACKNOWLEDGEMENTS

This work was supported by NIH grant GM56244 to ANI, by a Faculty Diversity Scholars Program award from the University of Massachusetts Medical School to TP-B, and by KAKENHI (JP17H03608, JP18H04802, JP19H05244, JP18H05527) and JST CREST JPMJCR16G1 to YO. We also thank the Advanced Computational Scientific Program of the Research Institute for Information Technology, Kyushu University.

## SUPPLEMENTARY DATA

**Supplementary Table S1.**
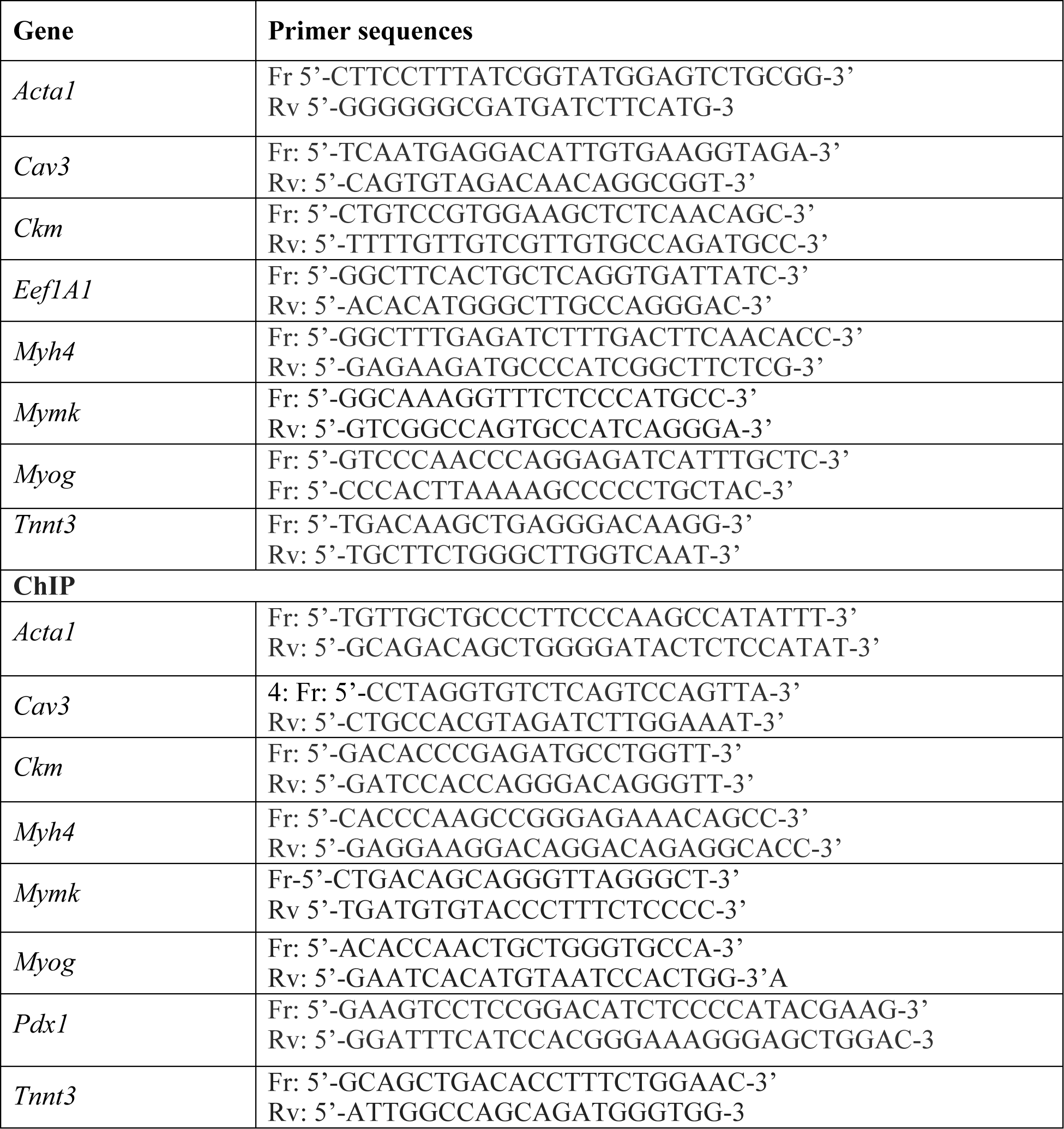
Sequences of forward and reverse primers used in Q-PCR

**Supplementary Table S2 (spreadsheet)**

RNA seq analysis: list of differentially expressed genes comparison DMSO vs FK506 at different time points

**Supplementary Table S3 (spreadsheet)**

RNA seq analysis: list of differentially expressed genes comparison different time points to GM conditions Clusters – list of genes

**Supplementary Figure S1.**
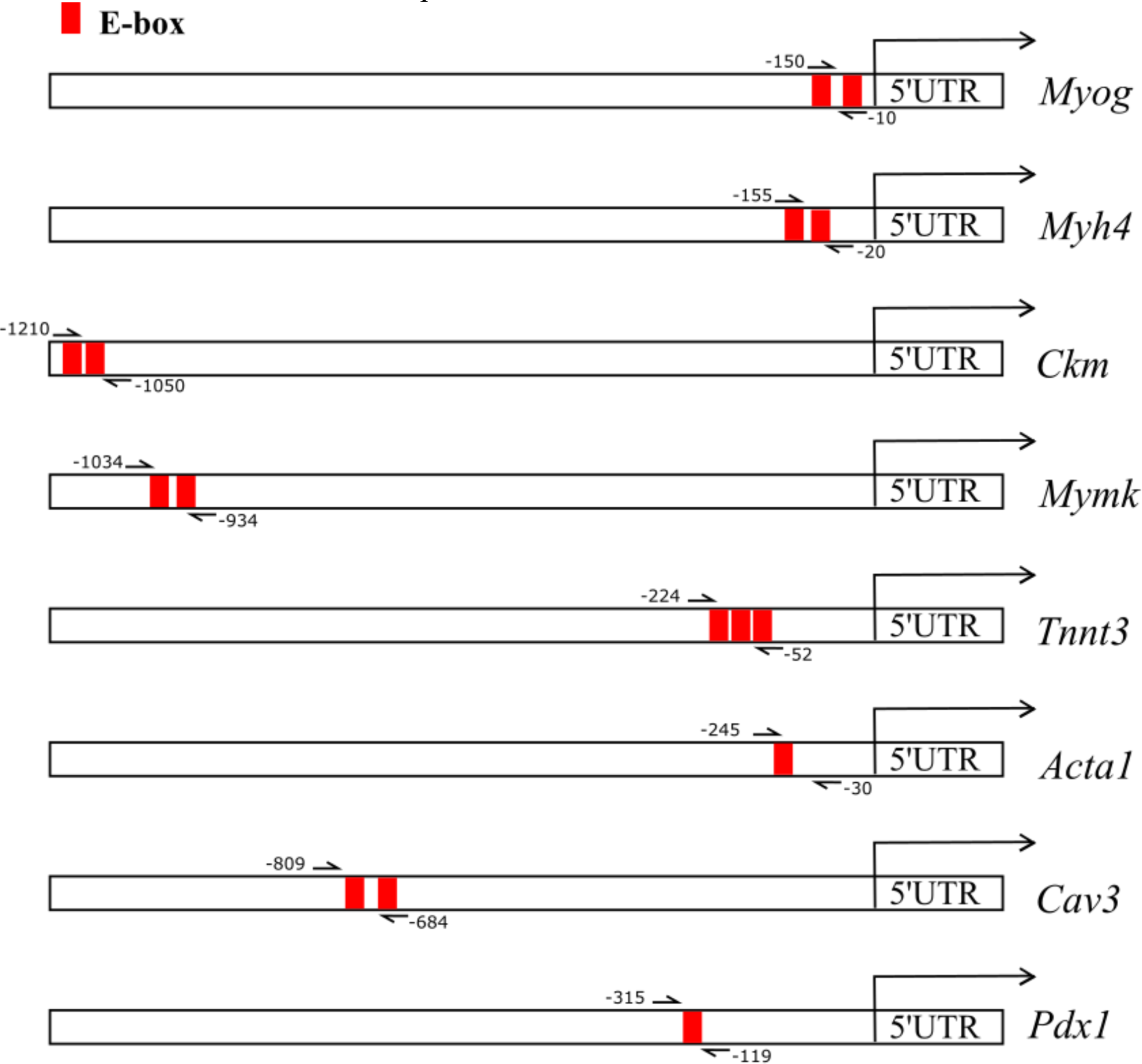
Schematic overview of the ChIP primers used.

**Supplementary Figure S2.**
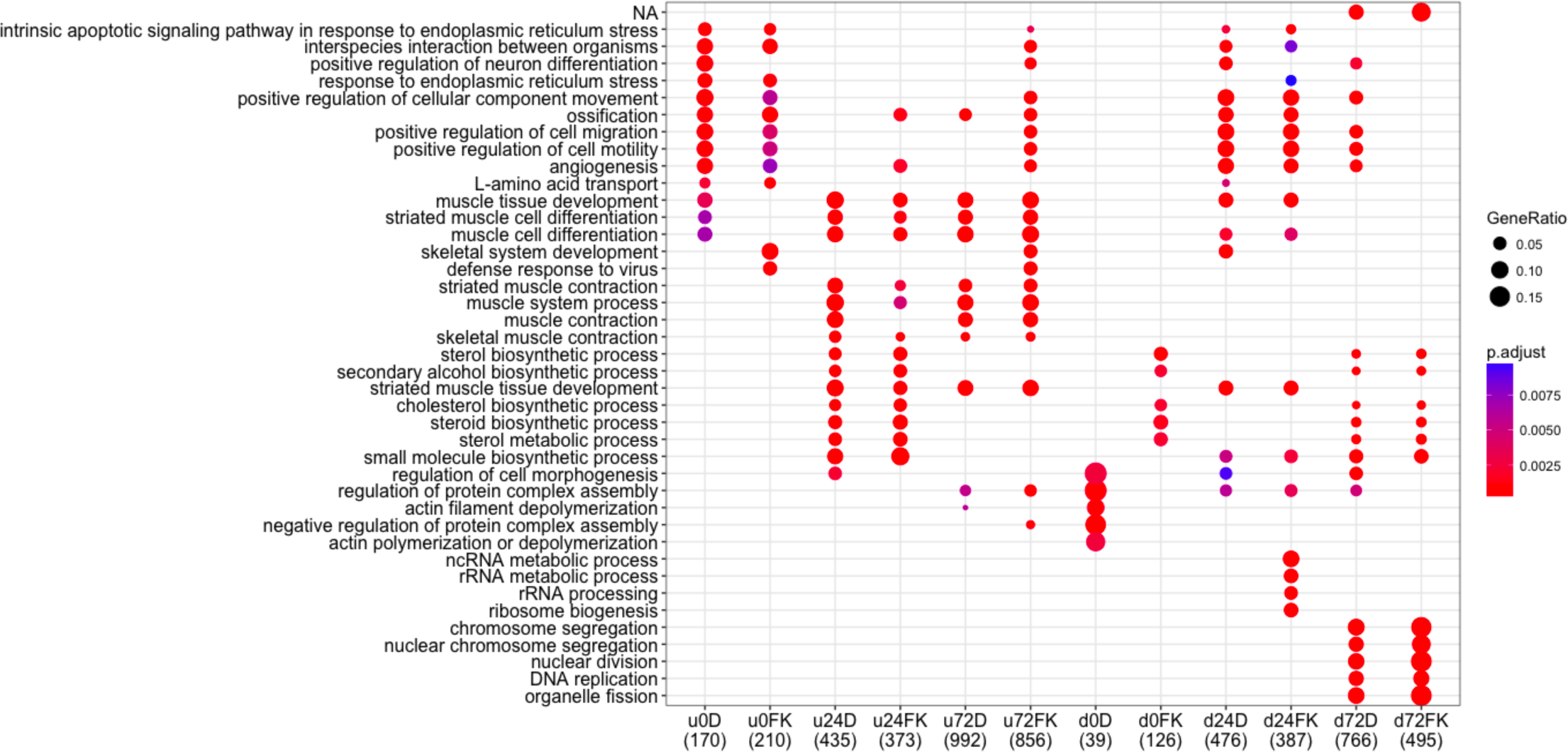
GO analysis of the biological processes shows differences between treatments (DMSO vs FK506) in muscle-related categories.

